# Enhanced 3D Osteogenic Differentiation of Encapsulated Human MSCs in Hyaluronic Acid Core Alginate Shell Capsules under Dynamic Culture Conditions

**DOI:** 10.64898/2026.09.29.755283

**Authors:** Julia Dreger, Daniela Andrea Contreras-Pérez, Maike Keck, Dominik Egger

## Abstract

The growing demand for three-dimensional (3D) in vitro models that accurately replicate human tissue to investigate disease mechanisms, enable tissue replacement, and support drug testing is driving the need for more advanced manufacturing platforms. Mesenchymal stem/stromal cells (MSCs) are promising candidates for tissue engineering and stem cell therapies due to their regenerative potential and ethical advantages. However, the reproducible and scalable production of 3D models that truly mimic the human complexity and function poses a major challenge. To address these challenges, we have developed a scalable and semi-automated 3D differentiation process for generating osteogenic microtissue derived from MSCs encapsulated in core-shell capsules (CSCs). To validate material-dependent, osteo-inductive effects of hyaluronic acid (HA), we systematically characterised our encapsulation process with regard to CSC integrity, spheroid formation and viability, as well as diffusion and shear thinning properties. Osteogenic differentiation was performed in static and dynamic culture, using either conventional 6-well plate format or a rotating wall vessel bioreactor. HA-CSCs were compared with inert carboxymethyl cellulose (CMC)-CSCs. In addition, hypoxic culture condition and the implementation of human platelet lysate (hPL) further supported physiological relevance. HA led to the formation of larger spheroids with high cell viability and increased metabolic activity compared to CMC. Calcium phosphate deposition within the extracellular matrix (ECM) and elevated ALP activity confirmed successful osteogenesis in all conditions. Furthermore, both HA and dynamic culture promote osteogenesis, and when combined, synergistically enhance the osteogenic differentiation. In conclusion, we present a semi-automated, high-throughput and xeno-free platform process for the 3D differentiation of MSCs towards osteogenic lineage. By integrating HA, hypoxia and dynamic culture conditions, this capsule-based system enhances the physiological relevance, enabling advanced in vitro disease modelling, drug testing, and scalable microtissue generation while reducing reliance on animal models.

Graphical Abstract

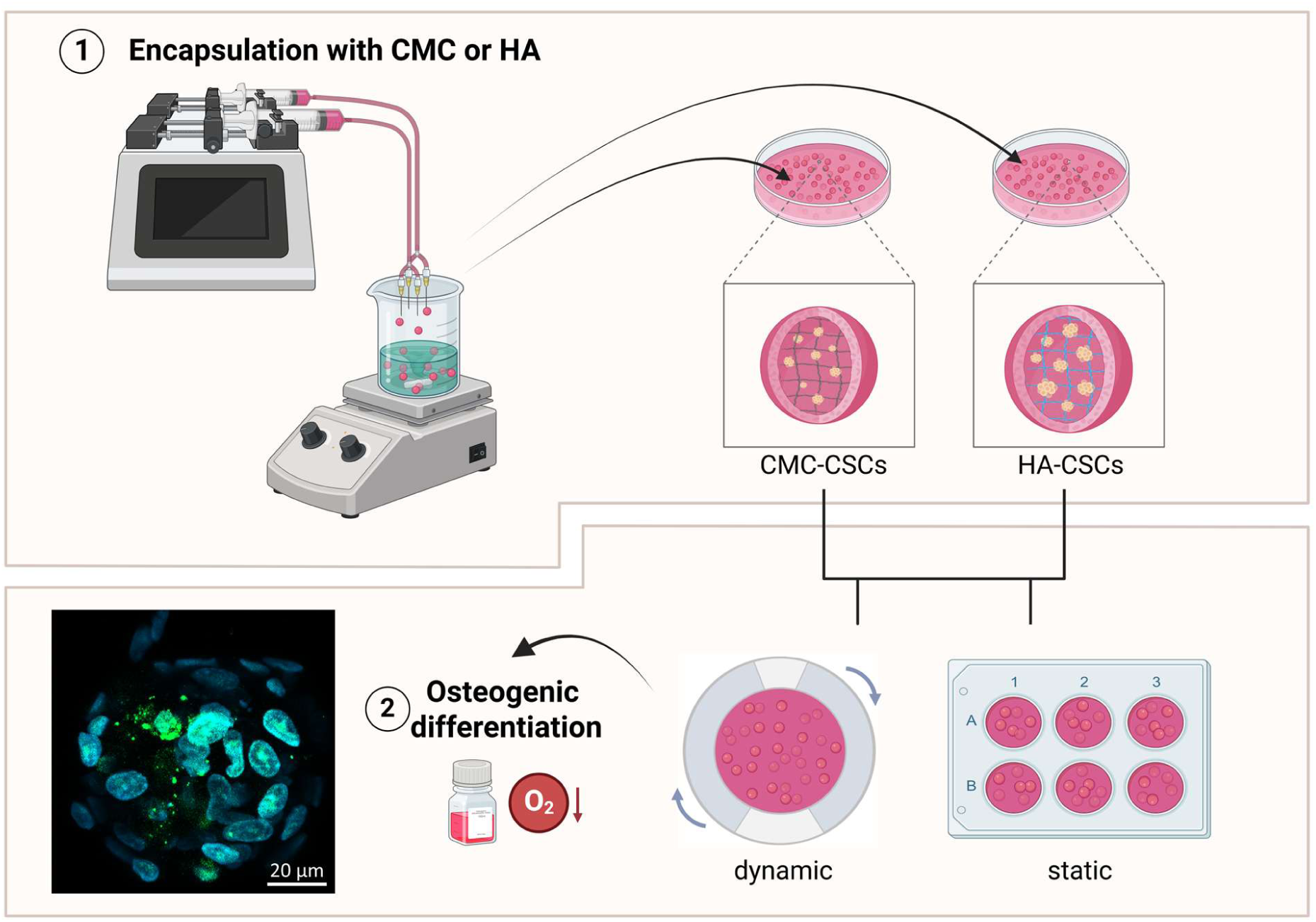

## 1. Introduction

Mesenchymal stem cells (MSCs) are widely used in tissue engineering due to their immunomodulatory properties and regenerative potential. MSCs can be isolated from many different tissue sources including adipose tissue, bone marrow or umbilical cord ^1,2^. Due to their ability to differentiate into multiple lineages, MSCs are a versatile source of cells for generating various tissue types, including bone ^3^. Osteogenic differentiation is highly relevant, since it represents an approach towards bone tissue engineering for larger and complex bone defects or to investigate skeletal diseases. Previous studies have shown that adipose tissue-derived MSCs (adMSCs) exhibit a notably high osteogenic differentiation potential compared to MSCs from other tissue sources ^4^. However, the differentiation potential of MSCs is highly sensitive to the applied culture conditions. A variety of factors, including oxygen concentration, nutrient availability and medium composition, influence cell behaviour and lineage commitment. To generate reliable and physiological data, it is therefore essential to mimic in vivo conditions as closely as possible ^5^. In regard to this, the use of reduced oxygen levels and xeno-free culture media is gaining more attention. Hypoxic culture conditions more accurately replicate the naive stem cell niche and have been shown to affect MSC proliferation and differentiation ^6–8^. Moreover, the use of human platelet lysate (hPL) supplemented media has been shown to increase cell proliferation and osteogenic lineage commitment compared to fetal bovine serum (FBS) supplemented media. Further, xeno-free media enhance translational relevance by eliminating animal-derived components ^9,10^. In addition, the choice of culture system is crucial. Evidence is growing that 3D cell culture systems demonstrate a more physiologically relevant approach compared to conventional static 2D culture, as they better mimic the in vivo microenvironment, which positively affects MSC stemness and lineage-specific commitment. Generally, 3D approaches can broadly be classified into scaffold-based and scaffold-free methods. Specially, scaffold-free spheroids have attracted considerable attention, as cell-cell interactions and spatial signalling are promoted and are associated with increased extracellular matrix (ECM) production, crucial for tissue formation and differentiation progression ^11–15^. Notably, Frith et al. (2010) and Wolff et al. (2023) demonstrated enhanced osteogenic differentiation potential in 3D MSC spheroid culture ^15,16^. Further, spheroid culture revealed altered gene expression resulting in upregulation of genes associated with hypoxia, inflammation or angiogenesis ^17^. Various strategies have been developed for spheroid generation, ranging from spontaneous self-assembly to collision-based methods. Common techniques range from static hanging drop culture, the use of low-attachment plates or microwell plates to dynamic systems including centrifugation, mixing in bioreactor platform ^18^. However, despite their widespread use, only a limited number of studies have established dynamic culture conditions for spheroid models. This is of particular interest since static culture systems show limitations in mass transfer which can lead to diffusion gradients of nutrients, oxygen or waste products. Through the dynamic environment, improved mass transfer efficiency leads to enhanced culture homogeneity, reduced diffusion limitations and a more physiologically relevant culture system. A common problem when implementing spheroids in a dynamic system is the high mechanical forces through shear stress. In turn, high shear forces can trigger cellular processes, influencing MSC stemness and differentiation potential. Furthermore, spheroids exhibit pronounced cell-cell adhesion and have been shown to undergo spontaneous fusion upon contact. Therefore, uncontrolled aggregation of several spheroids to multi-aggregates increase heterogeneity and reduce reproducibility ^19^. To overcome these limitations, alternative strategies such as core-shell capsule (CSCs) approaches have been developed. Liquid CSCs typically consist of two compartments, a liquid inner core that facilitates the self-formation of spheroids, and a surrounding semi-permeable shell that provides mechanical stability while maintaining sufficient mass transfer. Therefore, CSCs facilitate the formation of self-assembled, spatially separated spheroids, which can be used for dynamic culture while being protected from external shear stress ^20^. CSC-based approaches have already been successfully used to encapsulate various cell types, including MSCs, demonstrating their potential as a versatile platform for 3D cell culture ^21,22^. Further studies have focused on incorporating materials known to enhance stem cell properties or to promote differentiation. One often mentioned material in this context is hyaluronic acid (HA), a natural, hydrophilic glycosaminoglycan and component of the ECM. HA is known to influence cellular behaviour mainly through interactions with its receptor cluster of differentiation (CD) 44 which regulates cell – cell interactions and cell adhesion ^23^. Therefore, HA was also used as the core material in alginate CSCs by Park et al. (2020) who demonstrated high MSC viability and increased growth factor secretion resulting in enhanced angiogenesis ^24^. HA is particularly interesting in regard to osteogenesis as several studies indicate that HA exhibits osteo-inductive effects, thereby promoting the differentiation of MSCs towards osteoblasts ^25–28^.

To the best of our knowledge, there is currently no model that combines 3D osteogenic differentiation of MSCs within HA-containing CSCs while considering the aforementioned advanced cell culture methods. Therefore, we established a semi-automated and scalable process for the generation of HA-CSCs through inverse gelation (Figure 1). In this process, a single-cell suspension supplemented with calcium chloride (CaCl_2_) and HA is dropped into a stirred alginate bath, resulting in CSCs with a liquid HA containing core and a solid alginate shell. After a few washing steps and a second crosslinking in CaCl_2_ the CSCs can be used for subsequent experiments. Besides HA, we also performed encapsulation with the well-established and inert CMC, to evaluate the osteo-inductive effect of HA. Generated CSCs were first analysed regarding capsule integrity, properties and cell viability before conducting osteogenic differentiation. Osteogenesis was performed in static and dynamic culture using a 6-well plate and the CelVivo ClinoStar2 rotating wall vessel bioreactor, respectively. For assessment of ongoing osteogenic differentiation, osteogenic lineage-specific ECM staining was performed to visualise calcium phosphate deposition. Moreover, the activity of secreted alkaline phosphatase (ALP), an enzyme involved in bone mineralisation during osteoblast maturation, was evaluated. Both methods are well-established and commonly used in bone research ^29,30^.

**Figure 1:**
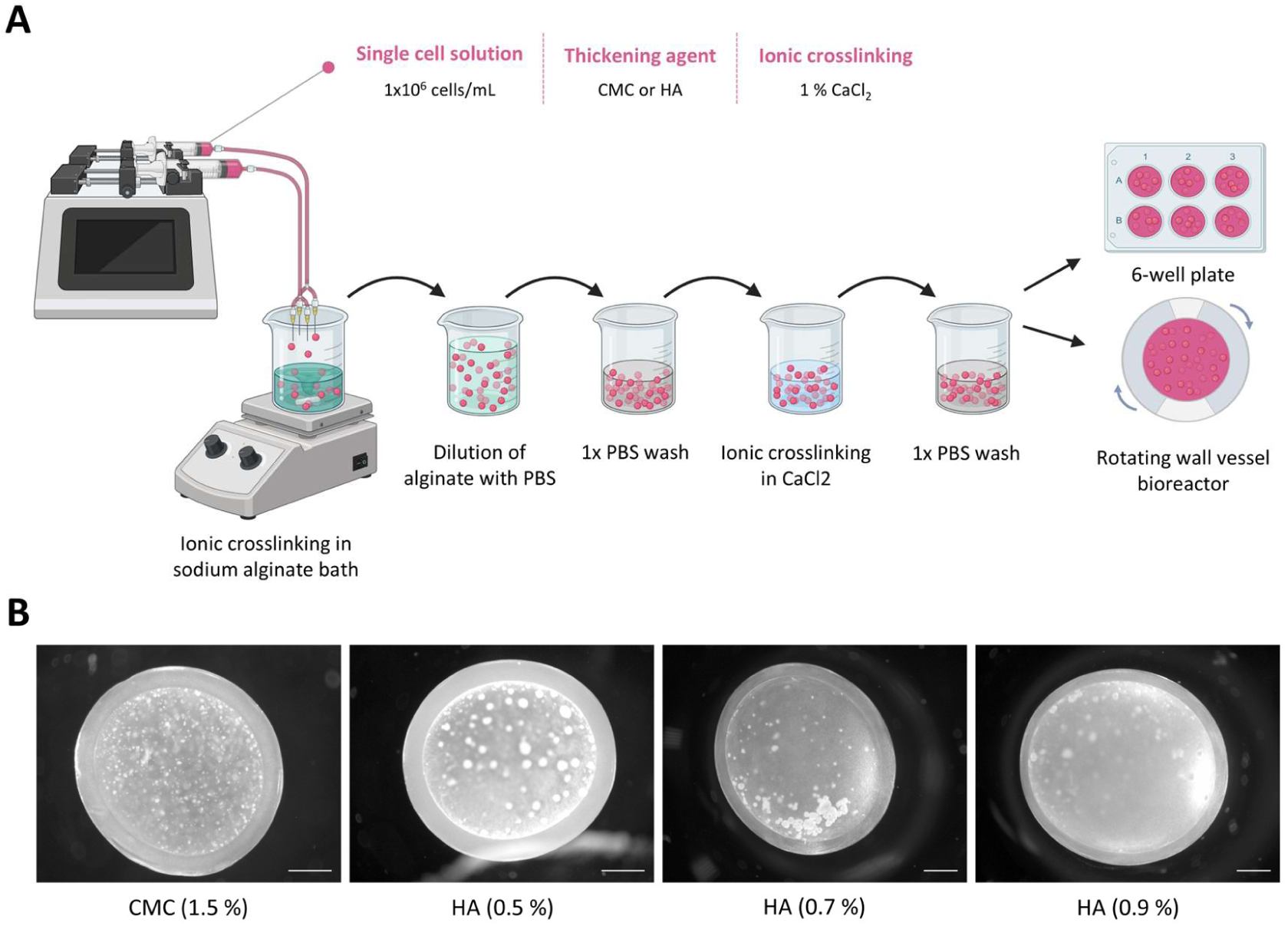
Encapsulation of MSCs in Core-Shell Capsules (CSCs). **(A)** Schematic representation of the encapsulation process using inverse gelation. Resulted CSCs can be transferred into different kinds of cultivation platforms e.g. 6-well plate or CelVivo ClinoStar bioreactor. **(B)** Representative darkfield images of CMC-CSC, 0.5 % HA-CSC, 0.7 % HA-CSC and 0.9 % HA-CSC, respectively. Scale bar: 500 µm.

## 2. Materials and Methods

If not stated otherwise all chemicals were purchased from Sigma Aldrich (USA).

### 2.1 Cell Culture

Human adMSCs were isolated from skin flaps removed during routine procedure. Isolation from human tissue was approved by the ethics committee of the University of Lübeck (reference number 20-233, November 2020 & 2024-455_1, November 2024). The patient gave written consent. The donor tissue was stored at 4 °C and processed within 24 hours of the surgical procedure. The adipose tissue was separated from the skin flaps, minced and digested with type IA collagenase for one hour. Following several washing steps, the stromal vascular fraction was transferred to a cell culture flask. The adMSCs were selected based on their adhesion to plastic and cell surface marker expression in accordance with the minimal criteria of the International Society for Cellular Therapy (ISCT). Unless otherwise specified, serum-free medium composed of alpha MEM (Thermo Fisher Scientific, United States), 2.5 % human platelet lysate (hPL; PL BioScience, Germany), and 0.5 % gentamycin was used for culture. Cells were cultured at 37 °C, 5 % CO_2_, and 5 % O_2_ in a humidified incubator to about 80 % confluence and detached with Accutase for harvesting. For cryo preservation, cells were suspended in cryo medium containing alpha MEM supplemented with 10 % hPL and 10 % dimethyl sulfoxide (DMSO) before long-term storage at −150 °C. All cells used for this project were expanded to a maximum of passage five.

### 2.2 Encapsulation of MSCs in Core-Shell Capsules

For CSC production, MSCs were harvested and suspended in one of the two thickening solutions containing culture medium supplemented with 1 % CaCl_2_ (Carl Roth, Germany) and either 1.5 % CMC or 0.5 %, 0.7 % or 0.9 % HA at a concentration of 1·10^6^ cells/mL. The cell suspension with CMC or HA was loaded into two syringes and connected to a tubing system. Through four 30 G cannulas, the respective solution was introduced into a 500 rpm stirred 0.5 % alginate bath at a flow rate of 1 mL/min using a syringe pump. After 5 minutes of inverse gelation, the alginate solution was diluted with the same volume of Phosphate Buffered Saline (PBS). Resulted CSCs were washed with PBS and stabilized through a second gelation step in a 1 % CaCl_2_ bath for 2 minutes. After two more washing steps, the CSCs could be collected and transferred into a cultivation platform. This involved both static cultivation in a 6-well plate and dynamic cultivation in a CelVivo ClinoStar bioreactor.

### 2.3 Characterization of CSC Dimensions

At least 15 capsules per condition were imaged using the Stemi 508 stereo microscope (Zeiss, Germany) and further analysed using ImageJ software. This enabled the determination of the outer and inner diameter, the shell thickness, and the sphericity factor of the CSCs as well as the diameter of the self-formed, encapsulated spheroids.

### 2.4 Determination of Diffusion Properties

The diffusion capacity was evaluated using a 4 kDa fluorescein isothiocyanate (FITC)-labelled dextran. Therefore, the cultivation medium was supplemented with 0.25 mg/mL dextran, in which CSCs were incubated. Over a 60-minute time series, fluorescence images were taken at 488/520 nm (Ex/Em) in 5-minute intervals by confocal laser-scanning microscopy (Eclipse TE2000-E, Nikon, Japan). To track the dextran enrichment in the capsule’s core, the bulk to core intensity ratio was calculated using ImageJ. Diffusion data were analysed using GraphPad Prism’s non-linear regression function to obtain values for the constant *k*. Following Fick’s law and assuming spherical shape leads to

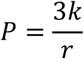

with *P* and *r* being the permeability coefficient and the inner radius of the capsule.

### 2.5 Rheological Characterization of Thickening Solutions

The viscosity of the thickening solutions was measured using an MCR 302 Modular Rheometer (Anton Paar, Austria) with a PP40 setup (40 mm diameter). Increasing shear rates up to 1000 s^-1^ were applied within 7.37 minutes. Extracted data were analysed using the power-law model

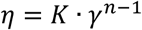

with 5 being the viscosity, *K* the consistency index, γ the shear rate and *n* the flow index. Logarithm leads to

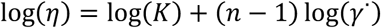

which was used to describe the behaviour of the fluids based on the slope. Each material was characterized in triplicate at 20 °C.

### 2.6 Evaluation of Cell Viability and Metabolic Activity

Cell viability was determined using Calcein acetoxymethyl (CalceinAM; Carl Roth, Germany) and Propidium iodide (PI) staining for visualising living and dead cells. Capsules were transferred into Eppendorf tubes containing 1 µL/mL CalceinAM and 3.3 µL/mL PI in PBS. After incubation for 30 minutes at 37 °C, CSCs were washed thrice with PBS and imaged using an Axio Observer A1 microscope (Zeiss, Germany). Furthermore, a resazurin-based in vitro Toxicology assay (TOX8) was used to evaluate the metabolic activity. For this, CSCs were transferred into Eppendorf tubes and weighed. In each case, 300 µL of TOX8 working solution was added per 100 mg of weight. After 3 hours of incubation at 37 °C, fluorescence intensity of the supernatant was measured at 560/590 nm (Ex/Em) using TE2000 Control 2.23.75 (Nikon, Japan). Data were blanked and values were normalized to day 0 capsules.

### 2.7 Recovery of Cells from CSCs

Encapsulated spheroids were harvested from capsules by dissolving the alginate shell chemically with sodium citrate. To achieve this, CSCs were collected in 15 mL low bind reaction tubes and dissolved in 100 mM sodium citrate solution (pH 7.4) for two minutes at room temperature. The sample was then diluted with five times the volume of culture medium and centrifuged at 500 × g for 5 minutes. The cell pellet was resuspended in PBS for further analysis.

### 2.8 Osteogenic Differentiation of Encapsulated MSCs

MSCs were expanded up to passage 4 and encapsulated in CSCs as described above. After 24 hours of spheroid formation, capsules were transferred into a 6-well plate (static cultivation) or into a CelVivo ClinoStar rotating wall vessel bioreactor (dynamic cultivation). The medium was replaced with osteogenic differentiation medium (alpha MEM supplemented with 0.1 μM dexamethasone, 2.5 % human platelet lysate, 0.5 % gentamicin, 0.2 mM L-ascorbic acid 2-phosphate and 5 mM beta-glycerolphosphate. Differentiation was performed in a humidified incubator at 37 °C, 5 % CO_2_ and 5 % O_2_ for a maximum of 21 days. Throughout differentiation, medium was replaced by half every two to three days.

### 2.9 Visualisation of Calcium Phosphate Deposition

For evaluation of differentiation, spheroids were harvested and stained on d0, d7, d14 and d21 for calcium phosphate deposition in the extracellular matrix. First, capsules were collected in 15 mL low bind reaction tubes before releasing spheroids with sodium citrate. In PBS suspended spheroids were centrifugated (500 × g, 5 minutes) and fixed with 70 % ethanol for 30 minutes at 4 °C. After centrifugation, spheroids were incubated in 5 µg/mL Calcein staining solution (Apollo Scientific, UK) overnight. Nuclei were stained in 5 µg/mL DAPI staining solution for 40 minutes and spheroids were subsequently analysed by confocal laser scanning microscopy (Eclipse TE2000-E, Nikon, Japan).

### 2.10 Quantification of ALP Activity

To determine alkaline phosphatase (ALP) activity during osteogenic differentiation, aliquots of culture medium were frozen during each medium change. Thawed samples were centrifugated at 14000 × g for 5 minutes. For each sample 80 µL supernatant was pipetted into wells of a 96-well plate prior to adding 20 µL of p-nitrophenyl phosphate stock solution (SIGMAFAST^TM^ pNPP and Tris buffer (Carl Roth, Germany) tablet dissolved in 4 mL ddH_2_0) and incubation at 37 °C for 40 minutes. Absorbance of the reaction product p-nitrophenolate was measured on a plate reader (Infinite 200 Pro M Plex, Tecan, Switzerland) directly after incubation at 405 nm). The ALP activity then is calculated via:

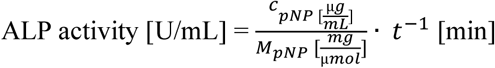

with *c_pNP_* being concentration of p-nitrophenolate, *M_pNP_* the molar mass of p-nitrophenolate and *t* the incubation time. Medium collected on day 0 served as blank control.

### 2.11 Statistical Analysis

For statistical analysis GraphPad Prism 10 was used (GraphPad Software, San Diego, CA, United States). Data was analysed using non-linear regression, one-way or two-way ANOVA tests assuming Gaussian distribution of residuals and equal standard deviation. Results were plotted and are presented as mean ± standard deviation (SD). Statistical significance is defined as: not significant P > 0.05; *P ≤ 0.05; **P ≤ 0.01; ***P ≤ 0.001; ****P ≤ 0.0001.

## 3. Results

### 3.1 CMC and HA 0.7 % Result in Different CSC Dimensions, However Still Exhibit Similar Viscosity and Permeability

To compare the effects of CMC and HA on capsule formation, generated CSCs were characterized by outer diameter, shell thickness and sphericity (Fig. 2A-C). The average diameters of CMC-CSCs and 0.5 % HA-CSCs were 2.42 ± 0.10 mm and 2.51 ± 0.09 mm, which were not statistically different from each other. CSCs with 0.7 % and 0.9 % HA turned out significantly larger (2.63 ± 0.13 mm and 2.85 ± 0.10 mm on average). While CMC-CSCs are the smallest, the shell is with 265 ± 31 µm the thickest compared to those produced with HA. Within the HA-CSCs the shell thickness is decreasing with higher HA concentrations (0.5 % HA: 246 ± 13 µm, 0.7 % HA: 191 ± 23 µm, 0.9 % HA: 180 ± 13 µm). Using CMC or HA resulted in capsules with a round instead of an elongated shape with the sphericity being below 0.1. Only capsules with 0.9 % HA had a significantly higher sphericity compared to the others.

**Figure 2:**
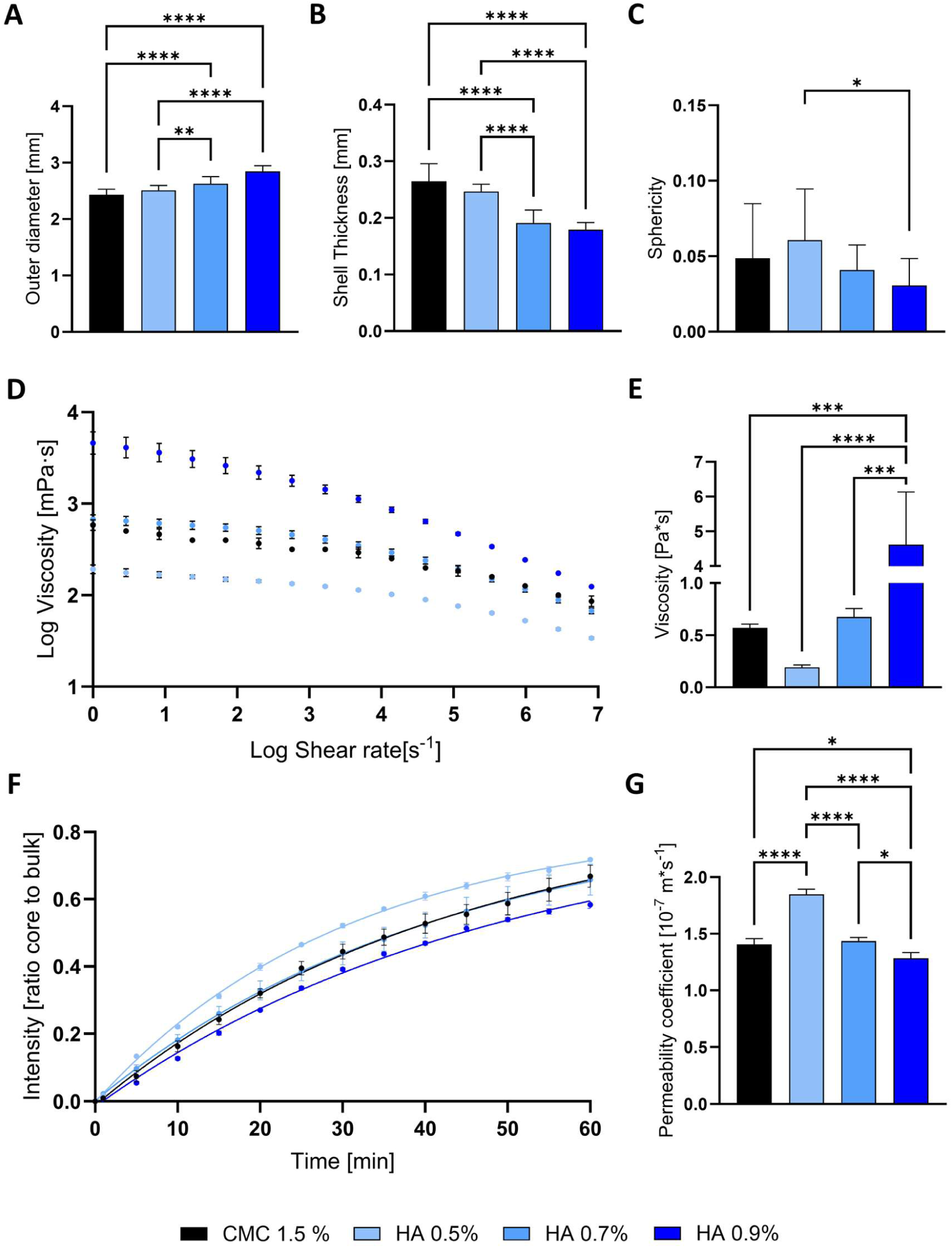
Characterization of capsule properties. **(A)** Outer diameter, (**B**) shell thickness and (**C**) sphericity of CMC- and HA-CSCs were assessed using ImageJ analysis on brightfield images; n ≥ 15. **(D)** Log viscosity as a function of log shear rate recorded via rheology measurement n = 3. (**E**) Viscosity of CMC and HA thickening solutions at two different shear rates, n = 3. **(F)** Saturation of 4 kDa FITC-labelled dextran in CSC’s cores; n = 3. (**G**) Calculated permeability coefficient from experimental data, n = 3. Data are represented as mean ± SD. Statistical analysis was performed using one-way ANOVA, n = 3 (*p ≤ 0.05; ****p≤ 0.0001).

Rheological measurements were used to evaluate the flow behaviour of the different core materials. To this end, the samples were each subjected to increasing shear rates. We found that the viscosity decreases when applying higher shear rates with *n* < 1, indicating non-Newtonian shear-thinning behaviour of all materials (Fig. 2D). In detail, *n*(CMC) = 0.89 ± 0.003, *n*(*HA*_0.5_) = 0.91 ± 0.005, *n*(*HA*_0.7_) = 0.88 ± 0.002 and *n*(*HA*_0.9_) = 0.81 ± 0.017 was determined. To elaborate further, viscosity at a shear rate of 1 s^-1^ was extracted (Fig. 2E). Results of the HA samples showed significantly reduced viscosity when decreasing the HA concentration (0.19 ± 0.02 Pa · s^−1^ at 0.5 %, 0.67 ± 0.08 Pa · s^−1^ at 0.7 %, and 4.62 ± 1.23 Pa · s^−1^ at 0.9 % HA). The similarity between HA 0.7 % and CMC (0.57± 0.03 Pa · s^−1^) can be corroborated by the extracted viscosity data showing no significant difference.

To determine the diffusion properties of the different capsules, diffusion of 4 kDa FITC-labelled dextran was reported. From this, the bulk to core intensity ratio was calculated and represented, showing saturation curves for both materials (Fig. 2F). Following statistical analysis using GraphPad Prism, all conditions reach a plateau at almost 80 %. In regard to this, analysis reveals a half-life of 30 minutes for CMC, 21 minutes for 0.5 % HA, 29 minutes for 0.7 % HA and 34 minutes for 0.9 % HA. Thus, CMC and 0.7 % HA exhibit similar dextran accumulation. Interestingly, CMC-CSCs and 0.7 % HA-CSCs exhibited similar permeability with 0.141 ± 0.005 µm/s and 0.144 ± 0.003 µm/s, respectively. In contrast, 0.5 % HA-CSCs revealed a significantly higher permeability coefficient compared to the other groups (0.185 ± 0.004 µm/s) while 0.9 % HA CSCs showed considerably lower permeability (0.128 ± 0.005 µm/s). Overall, out of all HA conditions, the diffusion and shear thinning properties of 0.7 % HA closest resembled those of CMC, while larger capsules were formed.

### 3.2 HA Promotes Formation of Larger, Viable Spheroids That Exhibit Extended Metabolic Activity Compared to CMC

Next, the influence of the different core materials on cellular level was examined. For this purpose, CSCs were produced with both materials and were cultured and examined over a period of five days. Encapsulated MSCs self-formed spheroids within the different core materials after one day (Fig. 3A). Interestingly, with 31 ± 9 µm on average, the diameter of the spheroids in the CMC core was significantly smaller compared to those formed in HA (Fig. 3B). Moreover, the HA concentration did not to have a significant influence on spheroid formation. In detail, capsules produced with 0.5 % HA resulted in spheroids with a diameter of 120 ± 44 µm, 0.7 % HA-CSCs in 117 ± 55 µm and 0.9 % HA-CSCs in 107 ± 31 µm spheroids on average. Differences could also be seen in the metabolic activity of the cells (Fig. 3C). TOX8 data indicates an approximately 4-fold increase for CMC-encapsulated MSCs within two days before activity decreases on day 3. In addition, metabolic activity increases in HA-encapsulated cells. Here, the activity increases progressively with higher HA concentrations (0.5 % HA: 3.1-fold increase on day 1, 0.7 % HA: 4.4-fold increase on day 2, and 0.9 % HA: 4.8-fold increase on day 1). Furthermore, at higher concentrations of 0.7 % and 0.9 %, HA exhibited prolonged and consistently high metabolic activity up to day four. This observation was neither found for cells encapsulated in 0.5 % HA-CSCs nor for cells encapsulated in CMC-CSCs. Interestingly, cells in CMC-CSCs and in 0.7 % HA-CSCs showed highest metabolic activity on d2 while d1 was peak for 0.5 % and 0.9 % HA.

**Figure 3:**
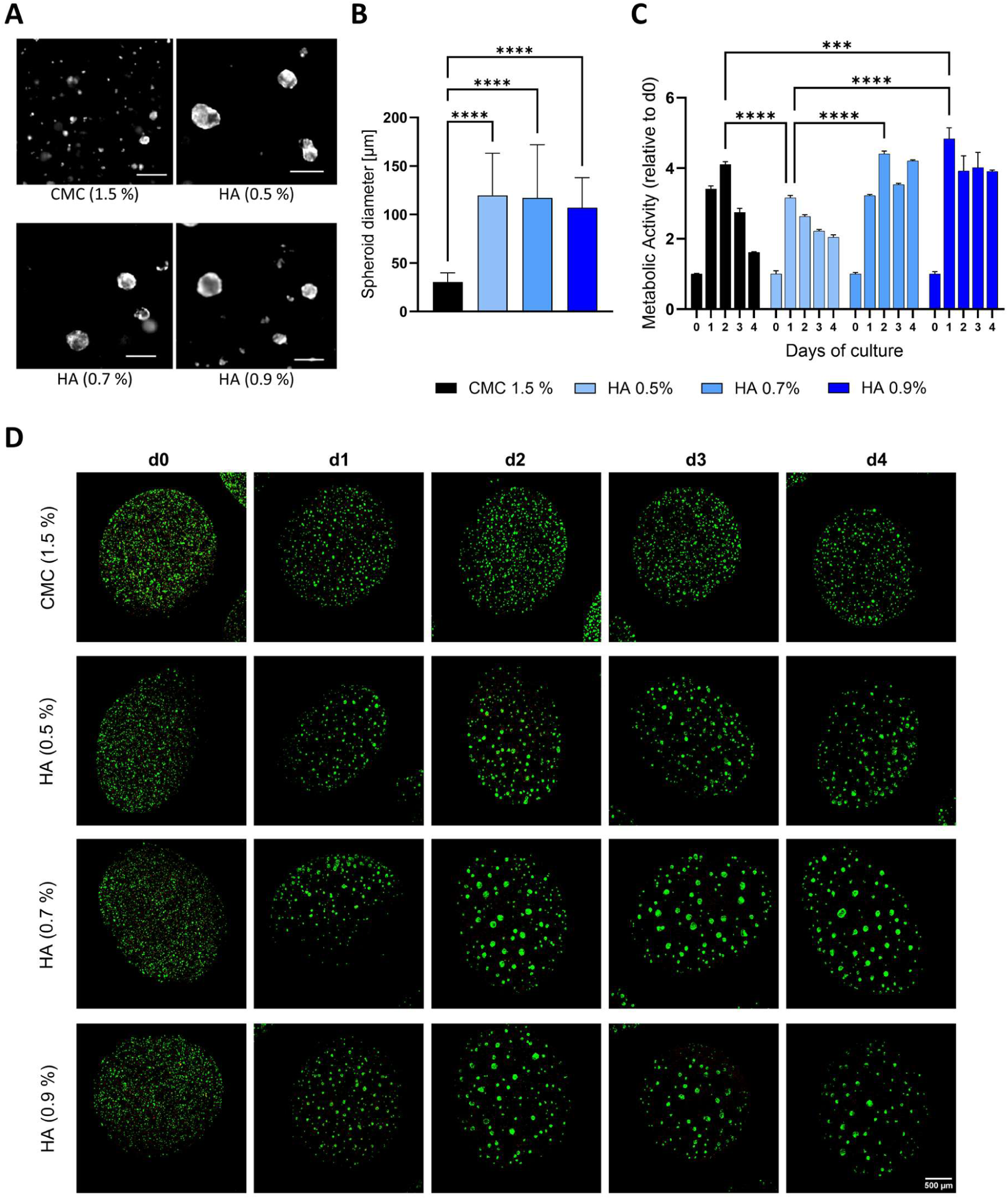
Influence of core composition on spheroid formation, cell viability and metabolic activity of encapsulated MSCs. **(A)** Representative fluorescence image of spheroid formation in CMC and HA-CSCs. Scale bar: 200 µm. **(B)** Spheroid diameter of encapsulated MSCs one day after encapsulation; n ≥ 15. **(C)** Resazurin based TOX8 assay for determination of the metabolic activity. TOX8 assay was performed on day 0-4 after encapsulation. Values were blanked and normalized to day 0 measurements; n ≥ 3. **(D)** Live/dead staining was performed on day 0, 1, 2, 3 and 4 of cultivation. Living cells are stained with CalceinAM (green) and dead cells with propidium iodide (red). Scale bar: 500 µm. Data are represented as mean ± SD. Statistical analysis was performed using two-way ANOVA (****p≤ 0.0001).

Finally, cell integrity was examined via live/dead staining and fluorescence microscopy immediately after encapsulation (day 0) until day 4 (Fig. 3D). High cell viability was demonstrated in CMC-CSCs during the first 4 days after encapsulation. Interestingly, even though the spheroids of HA-CSCs are 3-4-fold larger compared to those in CMC, high cell viability was also observed for HA in all conditions. In both materials, only a few dead cells were detected.

### 3.3 HA and Dynamic Culture Indicate a Synergistic Effect on Osteogenesis

Based on the results obtained so far, 0.7 % HA was selected for the subsequent osteogenic differentiation. This is because 0.7 % HA exhibited comparable shear thinning and diffusion properties as well as metabolic activity to those of CMC, while the other properties were not concentration-dependent. This facilitates to draw reliable conclusions regarding the osteo-inductive effects of HA, ensuring that any observations are attributed to cell – material interactions rather than to physical material properties like viscosity. In addition, the impact of static (6-well plate) and dynamic culture (rotating wall vessel, CelVivo ClinoStar) on osteogenesis was examined.

Osteogenic differentiation of MSCs in CMC- and HA-CSCs was performed for 21 days. During differentiation, cell viability was assessed on day 0, 7, 14 and 21 (Fig. 4). Fluorescence images of live/dead staining showed dead cells in all conditions. Interestingly, in CMC-CSCs spheroid population seemed to be divided in dead and living spheroids whereas spheroids in HA-CSCs consisted of both, dead and viable cells. In particular, spheroids formed in HA showed a dead inner core surrounded by living cells. This finding could be observed from d7 until the end of differentiation on d21. When comparing static and dynamic culture, no clear differences were seen for CMC or HA.

**Figure 4:**
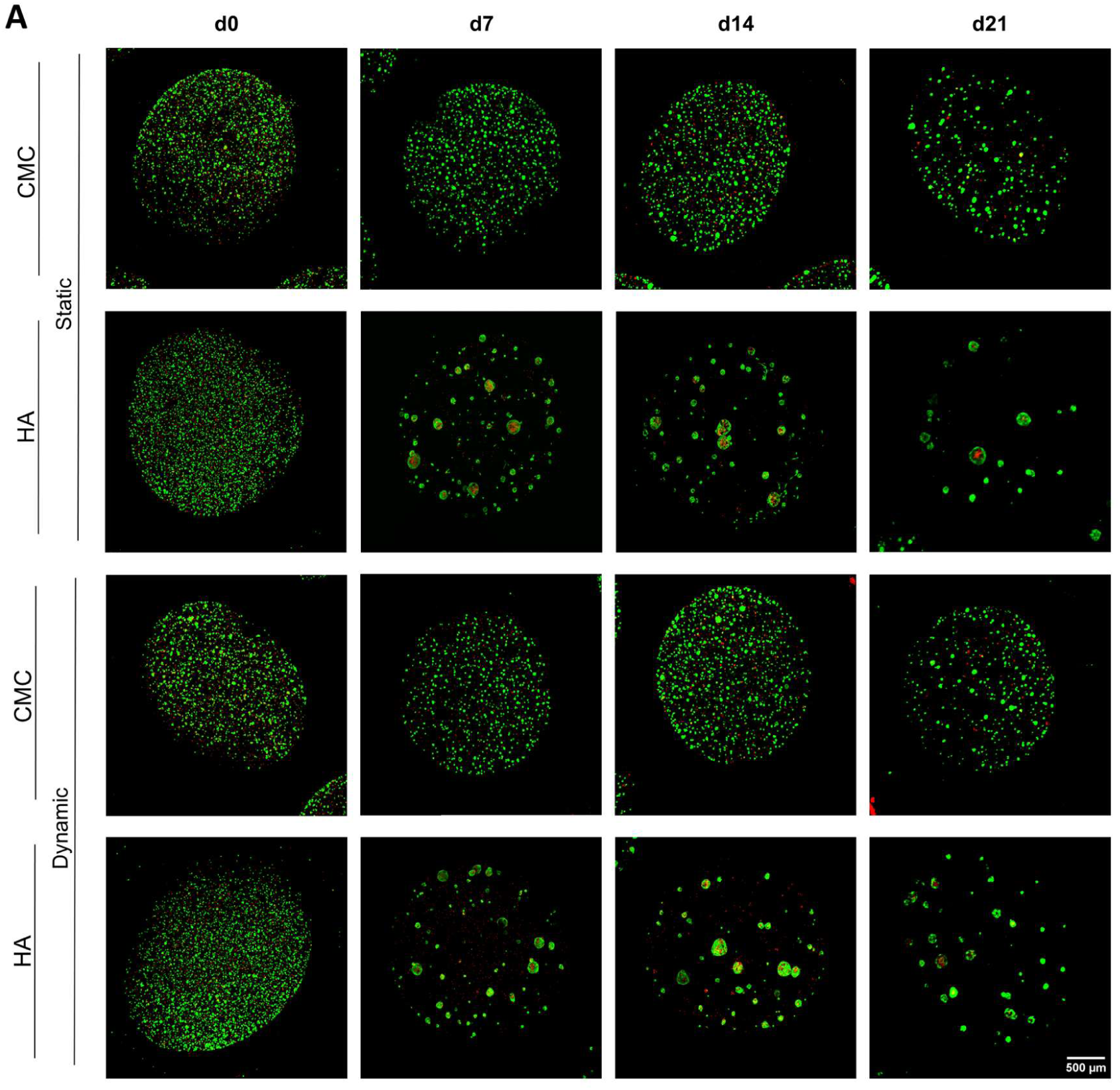
Sustained viability of CMC- and HA-encapsulated spheroids. Representative fluorescence images of CalceinAM/PI staining on day 0, 7, 14 and 21. Living cells were stained with CalceinAM (green) and dead cells with PI (red). Scale bar: 500 µm.

Osteogenesis was verified through Calcein staining and DAPI counterstaining, visualising calcium phosphate deposition and cell nuclei (Fig. 5A). The staining indicated that our capsule-based system facilitates osteogenic differentiation both in CMC and HA. In particular, ECM enrichment started from day 7 onwards and progressed until day 21. Initially, the Calcein signal was observed only in colocalization with the DAPI signal. As the differentiation progressed, extracellular accumulations became apparent. Following the differentiation, spheroids in HA-CSCs exhibited greater deposition compared to spheroids in CMC-CSCs. This effect was further amplified in HA-CSCs by dynamic cultivation, which resulted in stronger Calcein intensities. This finding was corroborated by the determination of the Calcein/DAPI ratio through semi-quantitative image analysis (Fig. 5B). During 21 days of differentiation, calcium phosphate deposition in the ECM was significantly increased in each condition (CMC static: 0.02 ± 0.02 to 0.71 ± 0.24; CMC dynamic: 0.02 ± 0.02 to 0.71 ± 0.24, HA static: 0.01 to 0.73 ± 0.29; HA dynamic: 0.04 ± 0.02 to 1.43 ± 0.56). Interestingly, we found that HA and dynamic culture, when considered separately, each led to a significantly higher calcium phosphate deposition compared to CMC-CSCs in static culture. The combination of HA and dynamic culture resulted in the highest ratio, which was significantly higher than each other condition, indicating a synergistic effect.

**Figure 5:**
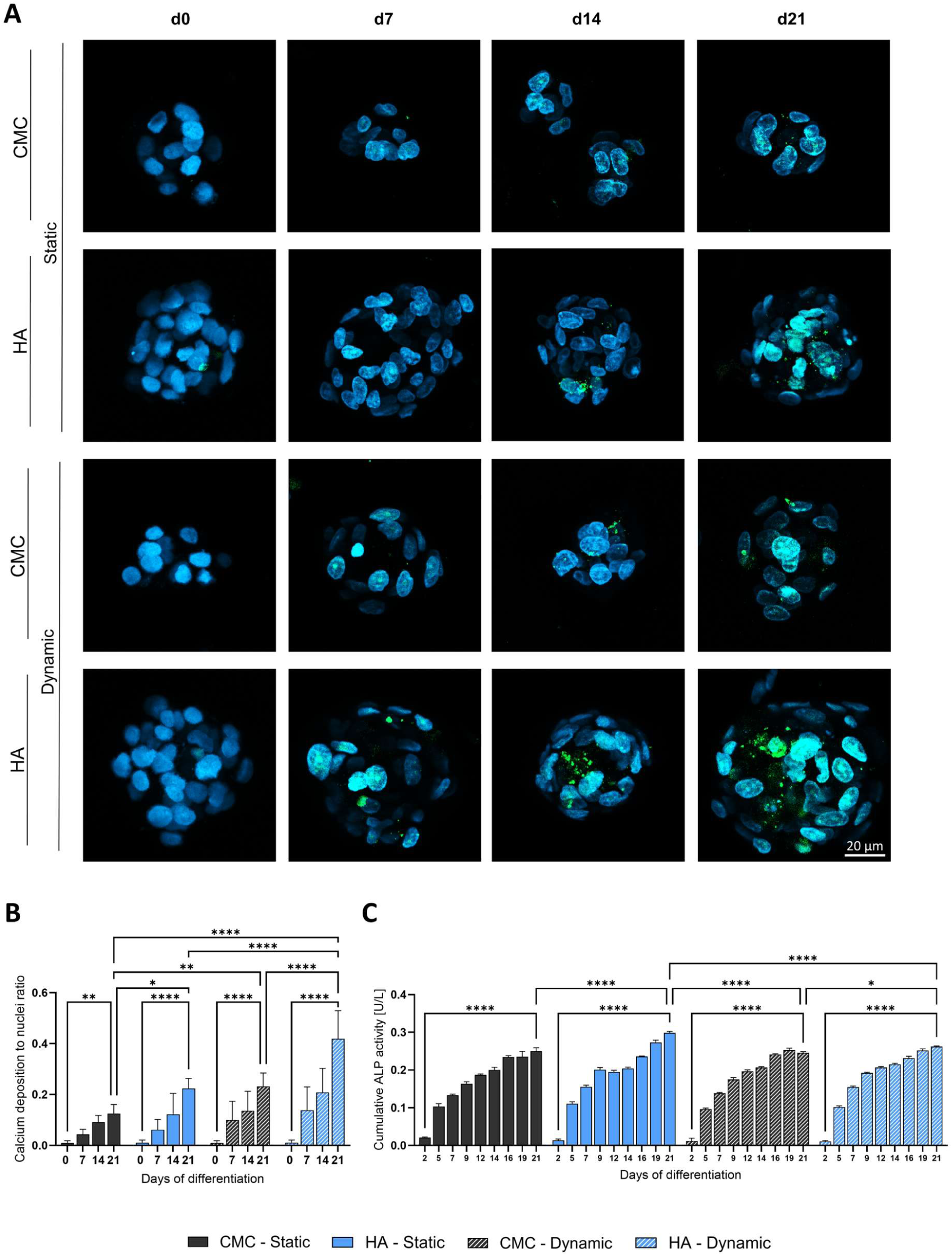
Osteogenesis potential of CMC- and HA-encapsulated spheroids. **(A**) Representative maximum intensity projections generated from fluorescence images using laser scanning confocal microscopy of Calcein (green) / DAPI (blue) staining on day 0, 7, 14 and 21. Step size: 1 µm; Scale bar: 20 µm. **(B)** Semi-quantitative image analysis of Calcein/DAPI stain using ImageJ software. Values are represented as area calcium deposition to area nuclei ratio, n ≥ 10. **(C)** Cumulative alkaline phosphatase (ALP) activity during 21 days of differentiation. Medium was collected after 2, 5, 7, 9, 12, 14, 16, 19 and 21 days; n = 3. Data are represented as mean ± SD. Statistical analysis was performed using two-way ANOVA (*p ≤ 0.05; **p ≤ 0.01; ****p≤ 0.0001).

Furthermore, we evaluated the ALP activity as an indicator of osteogenic differentiation (Fig. 5C). Increased ALP activity was demonstrated for both materials and culture conditions. With the exception of statically cultured HA-CSCs, ALP activity raised rapidly at the beginning of differentiation and followed a saturating trend towards d21. Here, no significant differences were found between those three conditions. In detail, ALP activity increased in statically cultured CMC-CSCs from 0.021 ± 0.002 U/L to 0.250 ± 0.009 U/L, in dynamically cultured CMC-CSCs from 0.012 U/L ± 0.007 U/L to 0.246 U/L ± 0.003 U/L and in dynamically cultured HA-CSCs from 0.011 ± 0.003 U/L to 0.262 ± 0.002 U/L. In contrast, ALP activity in statically cultured HA-CSCs increased similar at first, however reached a plateau from d9 to d14 before ALP activity increased again. After differentiation, an ALP activity of 0.298 ± 0.004 U/L was determined, which was significantly higher compared to the other conditions.

### 3.4 Static and Dynamic Culture Result in Differences Regarding the Dimension of Capsules and Spheroids

Lastly, the influence of the differentiation on capsule and spheroid dimension was examined. The results showed that the outer diameter of CMC- and HA-CSCs significantly increased during static culture (Fig. 6A). Compared to dynamic culture, the outer diameter decreased for CMC-CSCs and remained similar for HA-CSCs. In detail, the outer diameter changed from 3.13 ± 0.12 mm to 3.42 ± 0.18 mm and 3.16 ± 0.13 mm to 3.30 ± 0.16 mm in static cultured CMC- and HA-CSCs, respectively. For dynamic cultured CMC- and HA-CSCs, the diameter changed from 3.13 ± 0.12 mm to 2.81 ± 0.09 mm and 3.16 ± 0.13 mm to 3.06 ± 0.06 mm, respectively. Significant decrease in shell thickness was observed in all conditions besides static cultured CMC-CSCs (0.35 ± 0.03 mm before and 0.33 ± 0.05 mm after differentiation; Fig. 6B). In static cultured HA-CSCs, the shell was reduced from 0.39 ± 0.04 mm to 0.36 ± 0.01 mm. Furthermore, shell thickness decreased in dynamic cultured CMC- and HA-CSCs from 0.35 ± 0.03 mm to 0.30 ± 0.03 mm and from 0.39 ± 0.04 mm to 0.33± 0.03 mm, respectively. There were no significant differences in sphericity, all capsules retained their round shape (Fig. 6C). Finally, the influence on spheroid size was examined (Fig. 6D). Only spheroids in dynamically cultured HA-CSCs grew significantly from 86 ± 24 µm to 147 ± 35 µm during differentiation. All other conditions remained mostly the same size.

**Figure 6:**
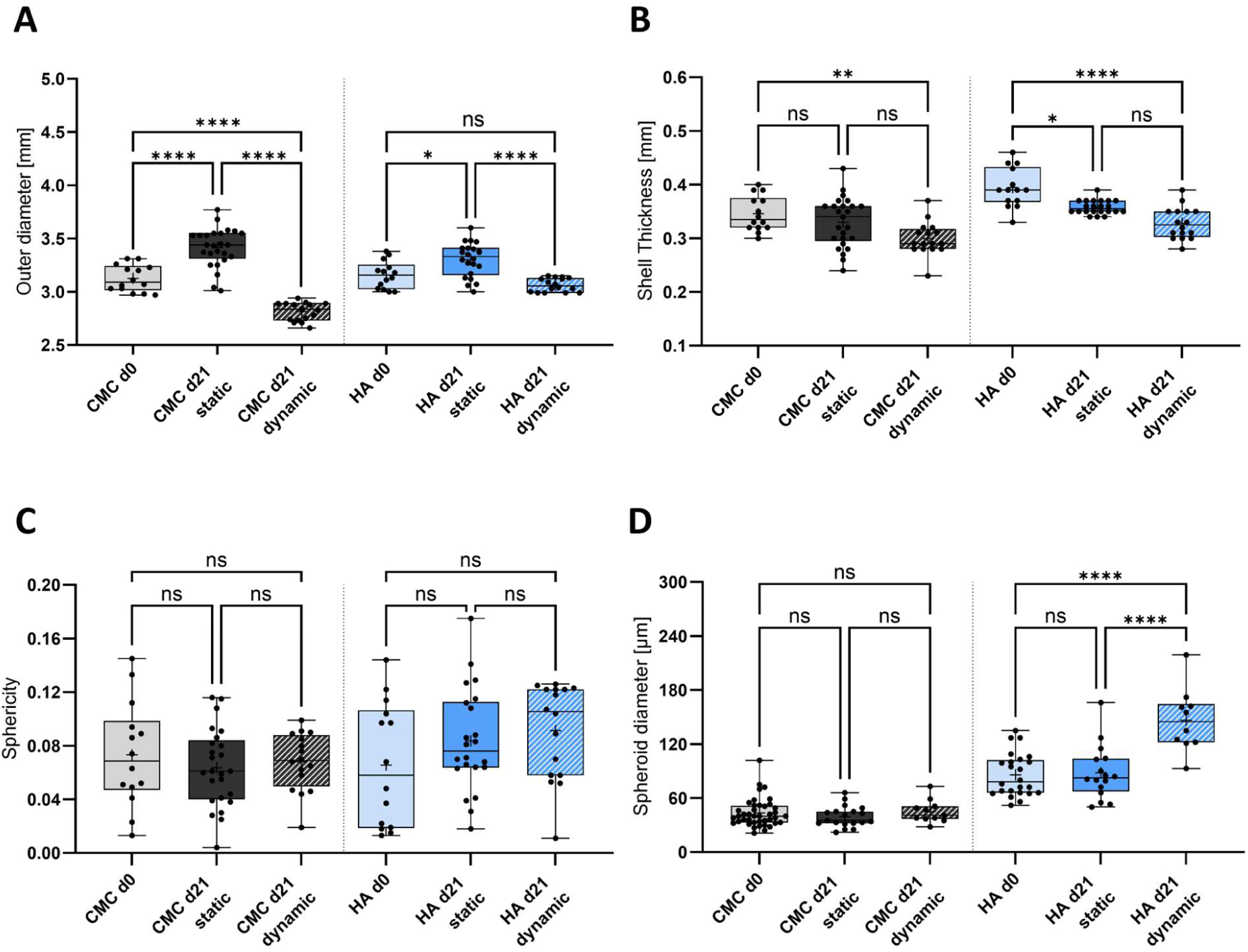
Comparative capsule and spheroid size analysis pre and post differentiation. Assessment of capsule’s (**A**) outer diameter, (**B**) shell thickness and (**C**) sphericity as well as (**D**) spheroid diameter on day 0 and day 21 of differentiation. Microscopic images were taken before and after differentiation and analysed via ImageJ software; n ≥ 10. Data are represented as mean ± SD. Statistical analysis was performed using one-way ANOVA (*p ≤ 0.05; **p ≤ 0.01; ****p≤ 0.0001).

## 4. Discussion

In this study, we described a semi-automated HA-based encapsulation process used for osteogenic differentiation of adMSCs. To the best of our knowledge, this is the first HA-capsule system used for osteogenesis of MSCs. Current studies focused either on osteogenesis in capsules but without HA ^31^, on osteo-inductive effects of HA in different 3D models ^27,28^ or used HA-capsules for the study of cell viability and angiogenesis ^24^. In addition, our system is distinguished by its use of xeno-free culture medium and the incorporation of hypoxia, further improving physiological relevance. As the aim of this study was to investigate the effect of HA on osteogenic differentiation, it was essential to minimise potential confounding physical influences. Therefore, the HA concentration was adjusted to match the viscosity and diffusion properties of CMC as closely as possible. This ensures that any differences observed between CMC and HA can be attributed to underlying cell – material interactions, rather than to physical system variations.

The observed increase in capsule size at higher HA concentrations may be due to the increased viscosity of the core solution. Rheological measurements confirmed that viscosity increases with higher HA concentrations. In particular, higher viscosity could affect droplet formation during the encapsulation process through altered surface tension on the cannulas resulting in larger droplets and consequently in larger CSCs ^32^. Despite the viscosity of 0.7 % HA being comparable to that of CMC, HA-CSCs were significantly larger. This suggests that material-specific properties of HA, in addition to viscosity, influence capsule formation. One possible explanation for this is the pronounced hydrophilicity of HA, which promotes the uptake of water and the subsequent swelling of the capsules ^23^. This hypothesis is supported by the observation of a thinner shell in HA-CSCs compared to CMC-CSCs. While the amount of CaCl_2_ within the HA-core remains constant, swelling increases the total core volume, thereby altering the surface to volume ratio. Consequently, the shell thickness of HA-CSCs decreases. Interestingly, despite 0.7 % HA and CMC-CSCs showed those geometric differences, the permeability of HA and CMC capsules was comparable. This might be explained by a compensatory effect whereby the larger core volume of HA is balanced by a thinner shell. This results in similar overall dextran transfer compared to diffusion in smaller CMC-CSCs with thicker shells.

Furthermore, encapsulated cells self-organized into spheroids within HA- and CMC-CSCs while remaining spatially separated from each other without formation of multi-aggregates. Notably, spheroids formed in CMC were significantly smaller compared to those in HA. This difference may be attributed to material-specific interactions between the cells and the HA matrix. HA is known to interact with cells through binding to the cell surface receptor CD44 ^23^. Interaction with CD44 can influence cell aggregation, proliferation and spheroid compaction, leading to the formation of larger spheroids. This assumption is supported by the observation that varying HA concentration showed no significant change in spheroid diameter. This further indicates a material-driven effect rather than a viscosity-dependent mechanism.

Hyaluronic acid seems to also influence the metabolic activity of the encapsulated MSCs. Similar to CMC, an increase in metabolic activity was observed for HA-CSCs. Notably, higher HA concentrations were found to result in a more pronounced increase than lower concentrations. Thus, 0.5 % HA and 0.9 % HA showed a 3- and 5-fold increase, respectively. In addition, metabolic activity remained elevated for a longer culture period at 0.7 % and 0.9 % HA compared to CMC. Particularly noteworthy is the comparison between CMC and 0.7 % HA. Under both conditions, metabolic activity reached its maximum on day 2, showing both an approximately 4-fold increase compared to day 0. While both conditions exhibited a similar initial increase, metabolic activity remained high until the end of the observation period on day 4 for MSCs encapsulated in HA-CSCs, whereas it decreased after day 2 for CMC-encapsulated MSCs. This behaviour may likewise be attributed to a material-driven property of HA. Since the spheroid size did not depend on HA concentration, it is unlikely that the increased metabolic activity resulted from the formation of larger spheroids. Instead, the observed increase in metabolic activity could be due to either enhanced cellular metabolism or an increased number of viable cells resulting from increased proliferation. Enhanced proliferation and increased metabolism induced by HA have previously been reported for both MSCs and periodontal ligament cells ^33–35^.

Live/dead staining revealed highly viable cells for both CMC and HA throughout the five days of culture. These findings align with previous reports by Kouroupis and Correa (2021), suggesting that spheroids with a diameter below 200 µm maintain high cell viability without elevated levels of cell death induced by diffusion limitations ^36^. In contrast, we were able to detect changes in cell viability during differentiation.

The observed decrease in cell viability during the three weeks of differentiation might be explained by increased compaction of spheroids and ECM production ^37^. Similar observations have been reported by Tatsumi et al. (2026), who suggest that controlled apoptosis in larger spheroids may promote ECM production and therefore enhance osteogenic differentiation ^38^. This could be further supported by our previous cell viability tests, where five days of static cultivation showed no evidence of changed cell viability (Fig. 3). However, during differentiation, a pronounced increase in PI signal was observed after seven days. This further underline that cell death might be associated with the onset of differentiation rather than prolonged culture. Another often described explanation for increased cell death is impaired mass transfer. Since dynamic culture conditions are known to enhance mass transfer, reduced cell death would be expected compared to static culture ^19,39,40^. However, no differences in cell viability were observed between those cultivation strategies, further indicating that limited mass transfer is unlikely to contribute to the decreased cell viability.

Assessment of osteogenic differentiation revealed a significantly accumulation of calcium phosphates regardless of the condition. The observation that, particularly at the onset of differentiation, Calcein initially localized intracellular suggests the formation of extracellular versicles (EVs), responsible for calcium phosphate transport. Earlier studies have shown that the process of bone mineralization can be divided into two distinct phases. First, matrix EVs are loaded with calcium and phosphate ions through the interaction of various proteins. These loaded EVs then pass through the cell membrane in the second phase, where the calcium phosphate deposits accumulate in the ECM and gradually form a network ^41–43^. As differentiation progressed, calcium phosphates were subsequently detected in the ECM of all conditions, supporting the findings of a two-phased bone mineralisation process. The accumulation was significant in HA and dynamic culture. Additionally, we were able to show that HA and dynamic culture significantly enhance the process of mineralisation. Moreover, combining both resulted in the greatest calcium phosphate deposition, indicating a synergistic interaction between those parameters. A similar effect has been previously described by Zhang et al. (2019) ^26^. Assessing ALP activity and calcium deposition, the authors found that HA alone has limited effects on osteogenic differentiation of human amniotic MSCs, however was significantly enhanced by combination with further osteogenic stimuli. For instance, bioreactor systems like rotating wall vessel constructs not only improve nutrient and gas transfer but also provide mechanical stimulation which can further enhance MSC differentiation towards bone tissue ^44,45^.

In contrast to the synergistic effects of HA and dynamic culture we observed for calcium deposition, this could not be replicated in ALP activity. Instead, an increase in ALP activity followed by saturation was observed under all conditions. This may reflect the transient function of ALP as an early osteogenic marker as ALP expression reaches a biological maximum during osteogenesis, limiting its ability to distinguish between different stages of differentiation ^46^. Another explanation why ALP activity did not further increase might be also related to the structural changes within the spheroids. As differentiation progresses, increasing compaction and ECM mineralisation may limit the diffusion of ALP. This effect might be further relevant due to the CD44 receptor-dependent regulation of spheroid aggregation and compaction upon HA binding. Consequently, measured ALP activity in the supernatant might underestimate the actual cellular activity^4,23,47^.

Size characterization before and after differentiation revealed similarities between capsules with HA and CMC. Both capsule types exhibited an increased outer diameter under static culture during the three-week differentiation, which might be due to capsule swelling through osmotic water uptake ^48^. Interestingly, this phenomenon was more pronounced in CMC- than in HA-CSCs. This might be due to the aforementioned water absorption of HA during the encapsulation process, thereby limiting further swelling during differentiation ^23^. In contrast, an opposite trend was observed for dynamic culture, with the CSC’s outer diameter being significantly reduced for CMC and HA after differentiation. This observation may be attributed to mechanical abrasion induced by CSC movement within the bioreactor. The rotational motion facilitates repeated capsule – capsule interactions, generating friction and shear forces that might lead to surface erosion and progressive material loss ^48^. This hypothesis is corroborated by the finding that the shell thickness was also significantly reduced under dynamic condition. Nevertheless, capsules remained structurally intact, demonstrating ensured long-term differentiation up to three weeks in a bioreactor environment. Lastly, spheroids within dynamically cultured HA-CSCs were the only ones that became significantly larger during differentiation. Previous studies showed similar behaviour with increased spheroid diameter during trilineage differentiation of MSCs, which has been associated with enhanced matrix deposition during differentiation ^4^. In line with this, our observation may further emphasise the enhanced osteo-inductive effect when combining HA with dynamic culture in a bioreactor system.

Overall, the presented results provide valuable insights into the properties of CMC and HA and their influence on osteogenesis under static and dynamic culture conditions. To further corroborate these findings and enhance the model’s translational relevance further studies are needed. In particular, increasing the number of donors and investigating different MSC-tissue sources would give more in-depth understanding of biological variability. In addition, investing the potential of HA and dynamic culture on trilineage differentiation of MSCs would facilitate broader applications in the field of tissue engineering ^4,49^.

## 5. Conclusion

In this presented study we demonstrated a 3D differentiation platform for the scalable osteogenic differentiation of MSCs in HA-CSCs under well-defined conditions. The impact of the biomaterials HA and CMC on capsule and spheroid formation and cell viability was thoroughly analysed and coordinated to minimise confounding factors and enable direct comparison of the different core materials. Successful differentiation was assessed through spheroid diameter characterisation and further evaluated by osteogenic-specific calcium phosphate deposition in the ECM and by secretion analysis of ALP from media supernatant. Furthermore, capsule integrity was monitored before and after differentiation. Our data showed stable capsule generation with self-forming, viable spheroids in the capsule’s core for both biomaterials. Subsequent osteogenesis was significantly improved through the implementation of HA as a thickening agent and dynamic culture using a rotating wall vessel bioreactor. Together with the integration of hypoxic culture condition and xeno-free cultivation and differentiation media, this 3D capsule-based system supports the physiological recapitulation of in vivo phenotype. This facilitates the use for in vitro disease modelling and drug testing, or for high-throughput generation of microtissues for building blocks.

## Author Contributions

JD: designed and wrote the manuscript, researched literature, analysed data, designed figures, performed experiments, reviewed, and approved the manuscript. DC: researched literature, analysed data, and performed experiments, reviewed, and approved the manuscript. MK: provided resources, reviewed, and approved the manuscript. DE: Projekt administration, designed, reviewed, and approved the manuscript. All authors have read and agreed to publish the manuscript.

## Acknowledgments

Selected artwork was used from or adapted from images provided by BioRender.com. The authors would like to thank Antonina Lavrentieva and Domenic Schlauch for providing access and training on the rheometer.

## Abbreviations

2D: Two-dimensional; 3D: Three-dimensional; adMSC: Human adipose tissue-derived MSCs; ALP: Alkaline phosphatase; ANOVA: Analysis of variance; CaCl_2_: Calcium chloride; CalceinAM: Calcein acetoxymethyl; CD: Cluster of differentiation; CMC: Carboxymethyl cellulose; CSC: Core-shell capsule; DMSO: Dimethyl sulfoxide; ECM: Extracellular matrix; EV: Extracellular versicle; FBS: Fetal bovine serum; Fig.: Figure; FITC: Fluorescein isothiocyanate; HA: Hyaluronic acid; hPL: Human platelet lysate; ISCT: International society for cellular therapy; MSC: Mesenchymal stem/stromal cell; PBS: Phosphate-buffered saline; PI: Propidium iodide; SD: Standard deviation; TOX8: In vitro Toxicology Assay Kit

## Data Availability Statement

The data that support the findings of this study are available from the corresponding author upon reasonable request.

## Ethics Approval Statement

The studies involving human cells were reviewed and approved by the ethics committee of the University of Lübeck (reference number 20-233, November 2020 & 2024-455_1, November 2024). The studies were conducted in accordance with the local legislation and institutional requirements. The patient gave written consent.

